# Multimodal AI agents for capturing and sharing laboratory practice

**DOI:** 10.1101/2025.10.05.680425

**Authors:** Patricia Skowronek, Anant Nawalgaria, Matthias Mann

**Author notes:** Correspondence and material requests may be addressed to M.M.

## Abstract

We present a multimodal AI laboratory agent that captures and shares tacit experimental practice by linking written instructions with hands-on laboratory work through the analysis of video, speech, and text. While current AI tools have proven effective in literature analysis and code generation, they do not address the critical gap between documented knowledge and implicit lab practice. Our framework bridges this divide by integrating protocol generation directly from researcher-recorded videos, systematic detection of experimental errors, and evaluation of instrument readiness by comparing current performance against historical decisions. Evaluated in mass spectrometry-based proteomics, we demonstrate that the agent can capture and share practical expertise beyond conventional documentation and identify common mistakes, although domain-specific and spatial recognition should still be improved. This agentic approach enhances reproducibility and accessibility in proteomics and provides a generalizable model for other fields where complex, hands-on procedures dominate. This study lays the groundwork for community-driven, multimodal AI systems that augment rather than replace the rigor of scientific practice.

---

Much of a scientist’s expertise is learned through hands-on practice, not from manuals. This implicit knowledge - the subtle variations to a protocol or the instinct for troubleshooting - is critical in technique-intensive domains like our field of mass spectrometry (MS)-based proteomics, yet it is rarely documented. As chemist and philosopher Michael Polanyi observed, “we can know more than we can tell” (Polanyi, 1966). This challenge is amplified by the constant turnover in academic labs, weakening reproducibility and making cutting-edge science less accessible. Existing automation helps, but most systems are rigid and limited to liquid handling, offering little flexibility for high-pace research.

Generative AI promises to lower barriers to scientific discovery. Large language models can already review literature, analyze bioinformatics data, and even orchestrate multi-step research workflows - from generating hypotheses and writing code to analyzing results and drafting manuscripts (Yao *et al*, 2023; Lobentanzer *et al*, 2025; Liu *et al*, 2025; Miao *et al*, 2025; Schmidgall *et al*, 2025; Gottweis *et al*, 2025; Roohani *et al*, 2025; Swanson *et al*, 2025). In life science, this concept is beginning to extend into the physical lab, where AI has been used to program liquid-handling robots (Boiko *et al*, 2023).

Yet these tools still do not address the written vs. tacit knowledge divide - they do not support researchers in their hands-on laboratory tasks. Here, we demonstrate an AI agent that uses video to capture, share, and apply expert knowledge. By linking digital protocols with physical laboratory work, this system provides scalable, personalized guidance on complex procedures and offers a path toward more accessible, reproducible science.

To provide this guidance, we designed the AI agent to mimic an experienced colleague – able to answer questions, spot errors, and draw on years of practical know-how. The agents’ responses, adapted to current equipment conditions and researcher skill level, are delivered through a multi-agent AI framework that is accessed via a chat interface.

Our system is built using Google’s Agent Development Kit (ADK), Gemini 2.5 models accessed through Google Cloud and the Model Context Protocol (MCP) for tool integration. A main agent interprets a researcher’s request and delegates it to specialized sub-agents (Figure 1A).

**Figure 1:**
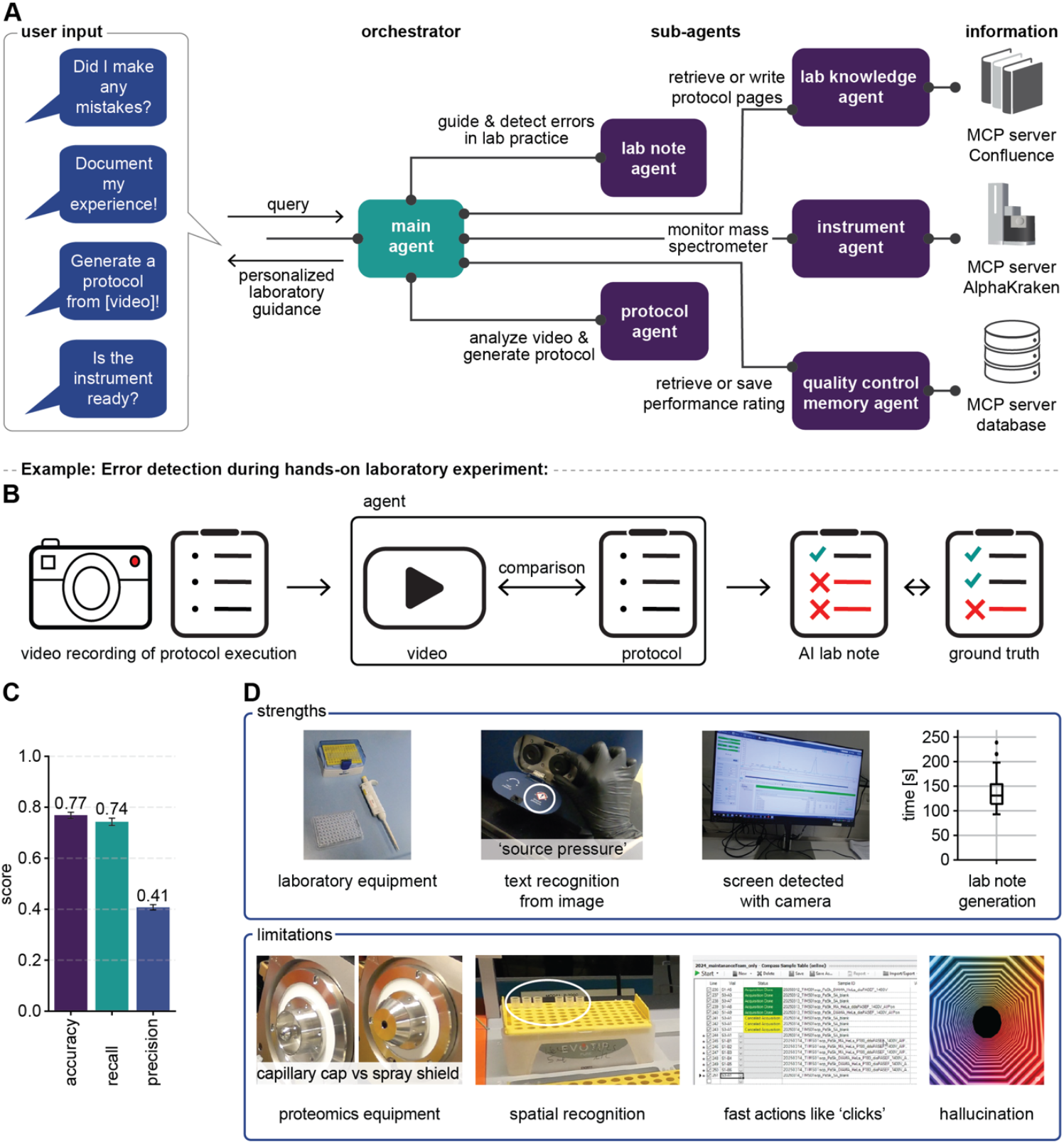
Proteomics lab agent for multimodal guidance. **(A)** The agentic framework translates conversational queries into personalized laboratory guidance. A main agent orchestrates specialized sub-agents: the Protocol Agent generates protocols from videos; the Lab Note Agent detects experimental errors; the Lab Knowledge Agent manages a knowledge base; the Instrument Agent monitors mass spectrometers; and the Quality Control Memory Agent accesses historical performance data. **(B)** Experimental workflow realized in our laboratory. Workflow where a scientist performed a protocol with intentional errors while recording the process. The AI agent analyzes the video against the baseline protocol to generate a lab note with flagged deviations. The agent’s performance was evaluated by comparing its AI lab note to a manually annotated lab note (ground-truth). **(C)** Overall agent performance metrics on the benchmark dataset, averaged over triplicate runs (error bars indicate standard deviation). **(D)** Examples of the agent’s strengths and limitations. Strengths include recognizing standard laboratory equipment, text on equipment, on-screen computer actions, and rapidly generating laboratory notes. Limitations include difficulty distinguishing visually similar, specialized proteomics equipment, achieving fine-grained spatial recognition (e.g., specific wells), detecting actions faster than 1Hz video sampling rate (e.g., mouse clicks), and occasional hallucination of events.

Each sub-agent has a specific function. For example, a Protocol Agent analyze a video of a person performing a specific sequence of actions to generate a protocol. Given an existing one, a Lab Note Agent detects errors or omissions when a researcher carries out the procedure. Other agents connect to the laboratory’s resources: the Lab Knowledge Agent retrieves documents from an internal knowledge base, the Instrument Agent monitors the performance of mass spectrometers, and a QC Memory Agent logs all quality control ratings to a central database, preserving the troubleshooting history for the entire team.

These agents are augmented with custom prompts encompassing a diverse set of examples and a proteomics knowledge base, enabling it to optimally reason through tasks. We evaluated the system’s performance after creating a benchmark dataset of lab protocols and videos (see Appendix).

To test the system in practice, we consider a common question when operating sophisticated analytical instruments: “Is this instrument ready for use?” To answer this, the agent goes beyond the latest quality control (QC) metrics, benchmarking current performance against historical data and expert decisions to issue a recommendation. It then records the researcher’s subsequent actions and supplies the necessary protocols for the next step (Appendix Figure S1).

This process contributed a continuously improving knowledge base, thereby building institutional memory that captures otherwise lost expert reasoning.

A key aspect of capturing expert knowledge is documenting procedures. Yet, writing informative, detailed and reproducible experimental protocols is a time-consuming process, which is often neglected. An AI can help in generating protocols using its expert knowledge but in our experience, this may involve invented details or hallucinations. Our agent avoids this by a video-to-protocol approach, grounding the process in direct observation. In practice, a researcher simply records their actions while explaining the procedures. Subsequently, the protocol agent analyzes the visual and audio data using its supplemented proteomics expertise, capturing the interactions with equipment, and generating a formatted protocol in minutes (Appendix Figure S2).

We compared the AI-generated protocols across ten diverse procedures, from simple pipetting to complex mass spectrometer operations to an established lab ground truth. The AI protocols achieved an average quality score of 4.5 out of 5 based on their completeness, technical accuracy, logical flow, safety instructions, and formatting (see Appendix & Appendix Figure S3). Overall, we found that AI-assisted protocol generation saves time while also creating detail-rich instructions that surpass the brief method sections of scientific papers.

Once the AI is equipped with a protocol it can in turn observe a researcher’s lab actions and point out omissions or mistakes. This should be especially helpful for researchers in training. To test this, we created a benchmark dataset by recording a scientist intentionally making 70 mistakes across 28 experiments (421 total steps). We then investigated whether our Lab Note agent could detect errors and omissions by comparing a researcher’s actions on video to a reference protocol and by flagging deviations (Figure 1B, Appendix Figure S4). Compared to ground truth, the agent correctly identified three-fourths of all mistakes, achieving an accuracy of 0.77 and a recall of 0.74 (Figure 1C). While precision was lower (0.41), indicating some false positives, a higher recall is preferable in this setting; it is better to flag a potential error for review than to miss a critical one.

Analyzing the AI error detection process, showed that the agent effectively identified general equipment and on-screen text, but struggled with tasks requiring fine-grained spatial recognition, domain-specific knowledge, and the detection of fast actions (Figure 1D, Appendix Figure S5). These limitations accounted for most of false positive identifications and explain why the agent successfully identified most omitted steps (91% recall) but missed subtle errors in execution (53% recall) or changes in step order (38% recall) (Appendix Figure S5D). Generating lab notes on average took just over two minutes and less than a dollar in token costs. These promising initial results can clearly be improved further as the abilities of multimodal agents advance fast, a trend we already experienced during development when upgrading to the more capable model Gemini 2.5 Pro, and can be further accelerated through domain-specific fine-tuning.

This study shows that AI can transfer practical expertise by linking written instructions with real laboratory work through multimodal understanding of video, speech, and text. The approach is adaptable to other scientific fields than proteomics that also rely on complex, hands-on procedures.

In the near term, such systems can provide real-time, interactive guidance that prevents errors before they happen and makes experiments more efficient. Over time, they can build an institutional memory that preserves expertise beyond individual researchers, standardize protocols, and even support predictive maintenance by tracking both instrument parameters and human actions. Once this system is fully implemented, scientists will literally be able to talk directly to these agents, receiving interactive guidance without having to stop and consult a written protocol.

This approach helps automate the routine parts of laboratory workflows that benefit from standardization, while preserving the human expertise needed for complex decisions. However, it is not a vision of blind automation in which tacit and artisanal knowledge is lost, but rather a way to capture and share it.

Realizing this potential to democratize science by spreading best practices requires a shared commitment. To this end, this study contributes a public benchmark dataset for evaluating AI on complex tasks in a high-barrier scientific domain. Its further development should be a community effort, embedding shared standards for transparency and supervision so that these tools augment rather than replace the rigor of scientific practice. If achieved, such AI partners could fundamentally transform laboratory work.

## Data availability

Complete code for agent and analysis is available on GitHub and the benchmark dataset consisting of videos, protocols and ground truth lab notes is available on Zenodo at https://doi.org/10.5281/zenodo.17253029.

## Supplementary Material

This manuscript contains an Appendix with additional figures and extended methods.

## Potential confiicts of interest

AN is an employee of Google. MM is an indirect investor in Evosep Biosystems. All other authors declare that they have no conflict of interest with the contents of this article.

## Acknowledgments

We thank our colleagues at the department of proteomics and signal transduction for their support and fruitful discussions. We are grateful for the input and constant support throughout the entire project from Magnus Schwörer and Fabian Opitz.

## Supplementary material for

This Appendix contains additional figures and extended methods.

**Appendix Figure S1:**
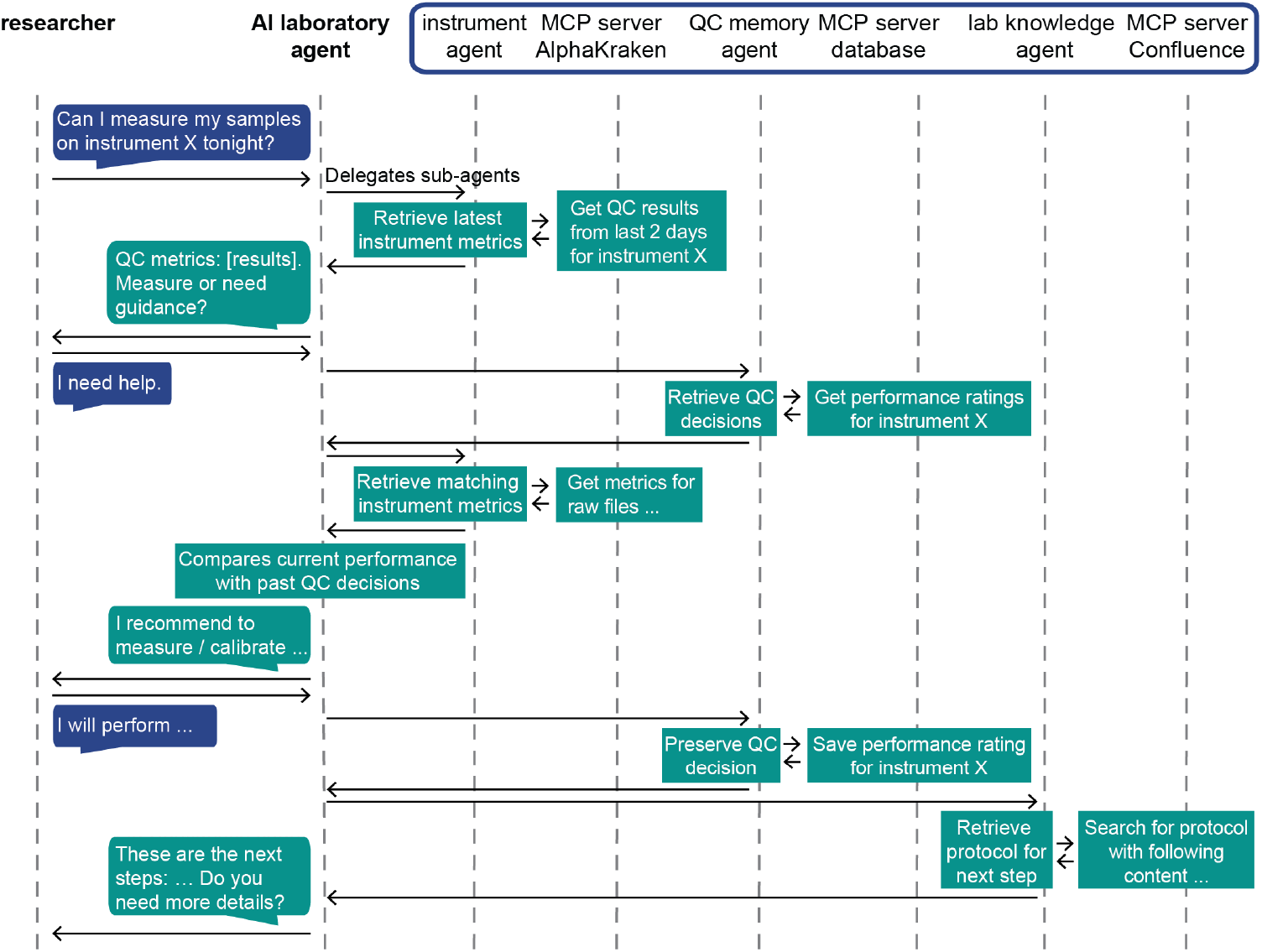
Agentic workflow for instrument performance queries. The flow of information between the user, the Main Agent, and the specialized sub-agents (Instrument, QC Memory, Lab Knowledge) during an instrument performance query.

**Appendix Figure S2:**
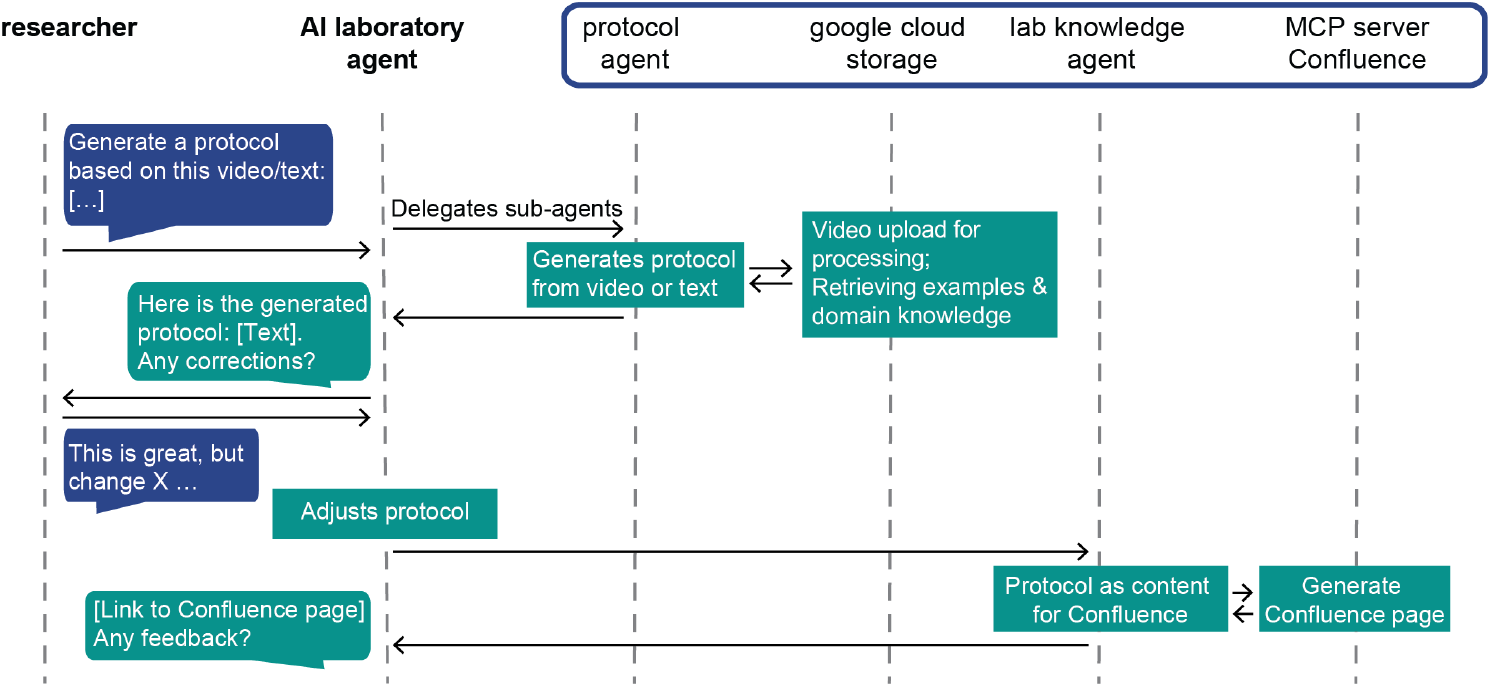
Agentic workflow for automatic protocol generation. The process from video upload to the final, user-approved protocol stored on a documentation platform such as Confluence.

**Appendix Figure S3:**
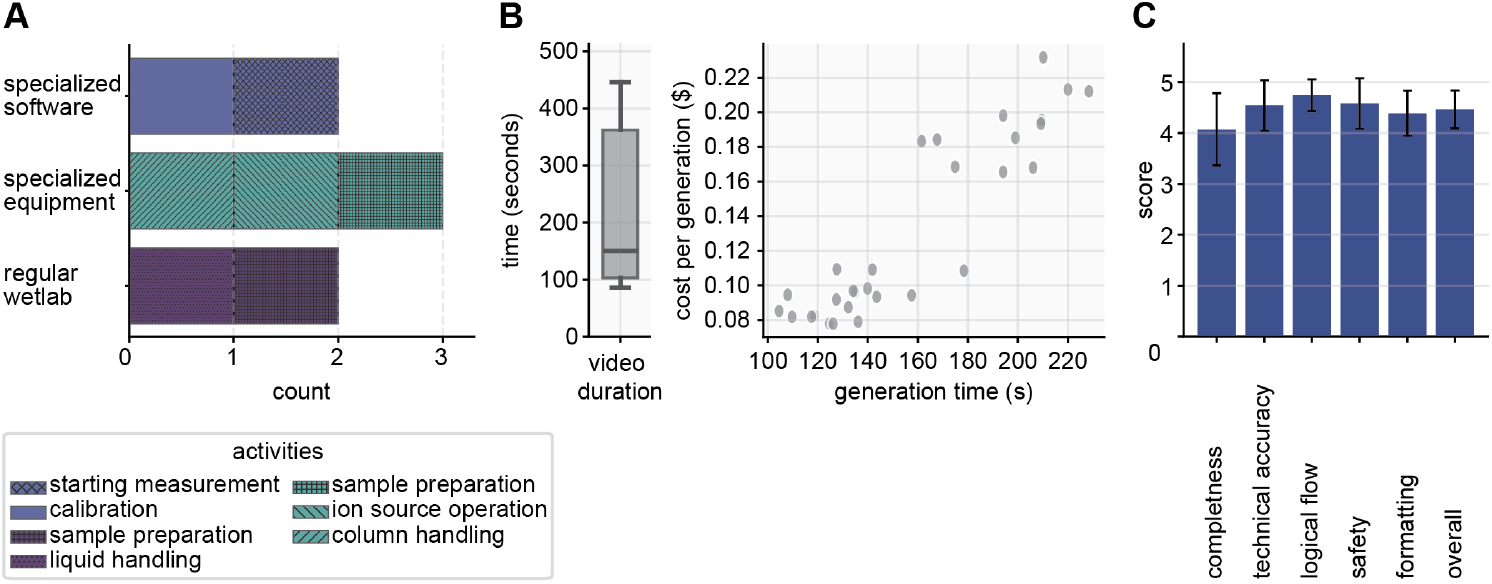
Performance metrics for the protocol generation task. **(A)** Composition of the benchmark dataset, categorized into specialized software, specialized equipment, and regular wet-lab protocols. **(B)** The left panel shows a box plot of the source video durations. The right scatter plot illustrates the relationship between the protocol generation time (in seconds) and the corresponding token cost per generation (in US dollars) for each protocol. **(C)** Evaluation scores for the generated protocols (in triplicate runs) across six categories: completeness, technical accuracy, logical flow, safety instructions, formatting, and an overall quality. The height of each bar represents the average score, and the error bars indicate the standard deviation.

**Appendix Figure S4:**
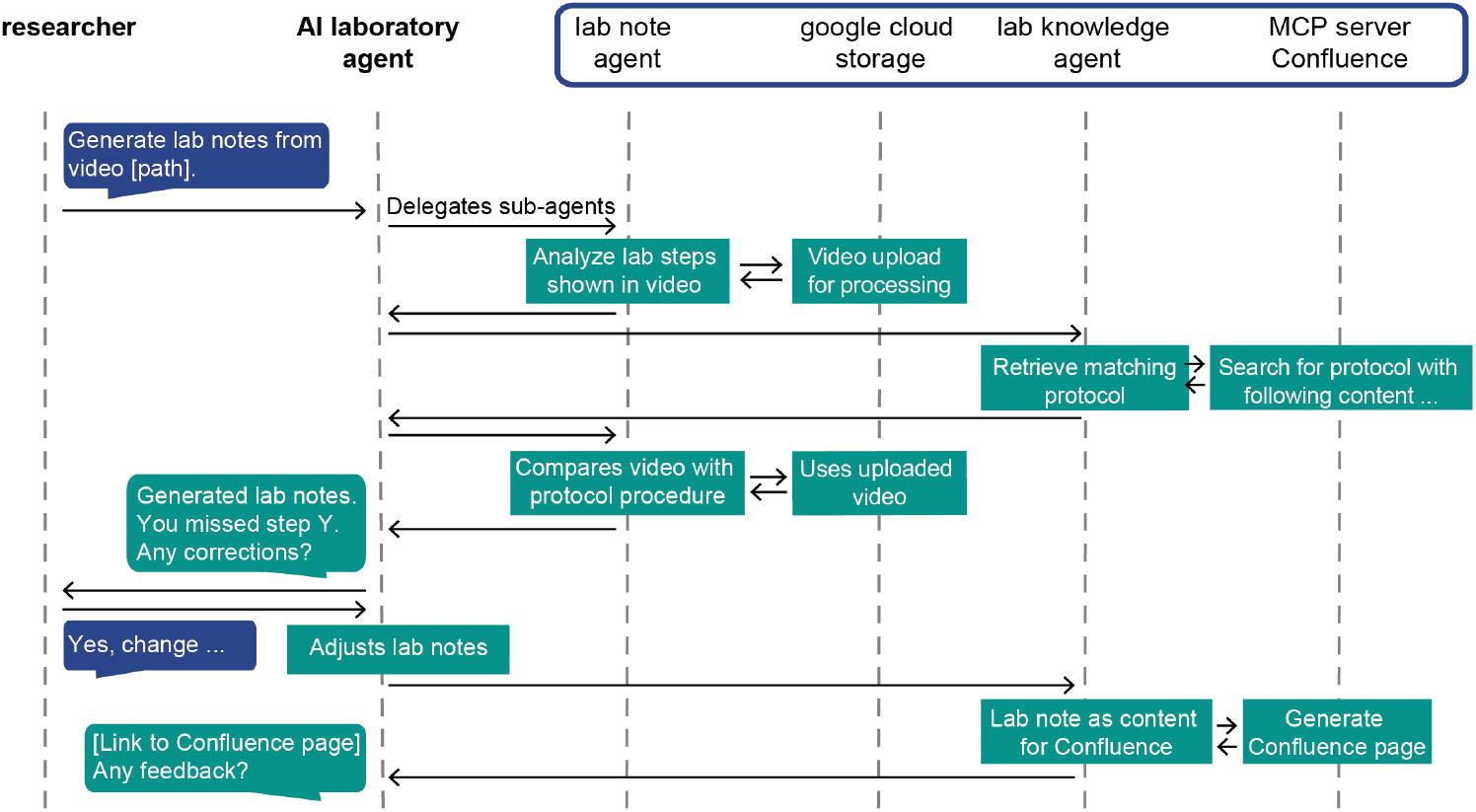
Agentic workflow for error detection and lab note generation. Illustration of the steps involved in identifying a protocol, comparing it against a video, flagging deviations, and generating final lab notes.

**Appendix Figure S5:**
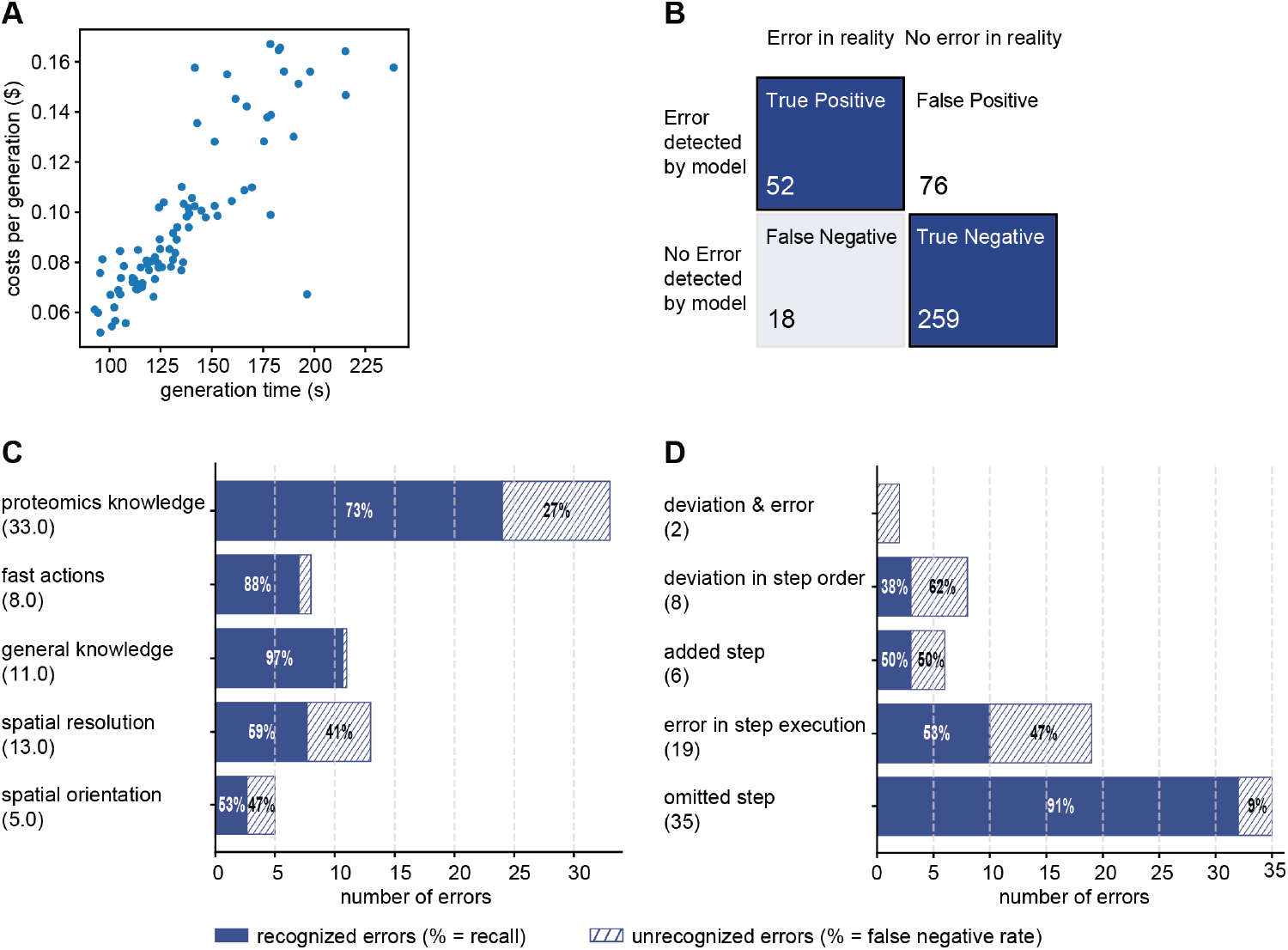
Performance of error detection and lab notes. **(A)** Relationship between the generation time and the associated token cost (in US dollars) for each lab note created. **(B)** Confusion matrix summarizing the model’s error detection performance on the benchmark dataset. **(C)** Errors categorized by the underlying AI skill limitation: Domain-specific proteomics knowledge, detection of fast actions, spatial resolution (e.g., reading the volume on a pipette) or spatial orientation (e.g., exact tip position in a rack). The numbers in parentheses indicate the total number of introduced errors for each category. **(D)** Error recognition by the type of protocol deviation.

### Extended methods

#### Agent framework

The system is a multi-agent AI framework designed to provide laboratory guidance and capture institutional knowledge. It is built on Google’s Agent Development Kit (ADK) and uses Google’s Gemini models as its foundation. Specifically, Gemini 2.5 Flash was used for the agent’s main functionalities, while Gemini 2.5 Pro was used for more complex video analysis. The agent accesses external data sources and tools such as Confluences pages via the Model Context Protocol (MCP). For video analysis tasks, a custom proteomics knowledge base and video files were retrieved from Google Cloud Storage and provided directly to the prompt.

All agent actions are directed by a Main Agent that functions as an orchestrator. It receives user requests from a chat interface, interprets them, and delegates tasks to a team of specialized sub-agents:

- Protocol Agent: Generates formatted, publication-ready protocols by analyzing video recordings of laboratory procedures (see below for more details).
- Lab Note Agent: Compares a video of a researcher executing a protocol against a baseline text protocol to detect errors, omissions, or other deviations. Subsequently, the agent generates a laboratory note flagging the found errors.
- Lab Knowledge Agent: Manages interactions with an internal knowledge base hosted on Confluence. In our implementation, the agent uses a local MCP server (mcp-atlassian) to search, retrieve, and generate Confluence pages.
- Instrument Agent: Retrieves current and historical performance metrics for mass spectrometers via a local [AlphaKraken](https://github.com/MannLabs/alphakraken) MCP server. AlphaKraken is an in-house developed tool for automated data processing and monitoring of mass spectrometry experiments.
- QC Memory Agent: Logs quality control (QC) decisions and user ratings to a local SQL database, accessed via an MCP server, to create a persistent history of instrument operation and troubleshooting.

#### Agent workflow

The role of the Main Agent is to be an AI research assistant with broad knowledge of proteomics. Its core logic matches user requests to predefined scenarios.

##### Scenario A: Instrument performance queries

This workflow is triggered when a user asks about instrument availability or performance (e.g., “Can I run my sample on tims1 tonight?”, where tims1 is one of the mass spectrometers). The agent follows a multi-step process (see Appendix Figure S1 above):

1. First the Main Agent invokes the Instrument Agent to retrieve the latest QC metrics from the AlphaKraken server.
2. The user is prompted to make a decision based on the data provided by the Agent (e.g., “Would you proceed with measuring or do you need guidance?”).
3. If guidance is requested, the QC Memory Agent retrieves historical decisions made under similar QC conditions (same instrument and analytical gradient) from the SQL database. This database stores the decision (measure: yes/no), a performance rating (1-5), and a comment linked to the specific raw file name, instrument and gradient. The remaining QC metrics of each raw file are again retrieved with the AlphaKraken MCP server.
4. The agent synthesizes the current and historical data to provide an actionable recommendation (e.g., “Other researchers measured at this performance level,” or “You should calibrate the TOF analyzer.”).
5. After the user decides on a course of action, the QC Memory Agent logs this decision, preserving it as institutional memory.
6. Finally, the Lab Knowledge Agent retrieves relevant protocols for the user’s next steps.

##### Scenario B: Automatic protocol generation

This workflow is triggered when a user requests to create a protocol from a video file or text notes (Appendix Figure S2). This requires a researcher to have recorded how they executed a procedure while explaining it, in order to create a tutorial video or notes about that procedure.

1. The Main Agent triggers the Protocol Agent, which uploads the video to Google Cloud Storage and then analyzes its visual and audio streams to produce a formatted, publication-ready protocol. Alternatively, one can use the notes as input.
2. The user can revise and approve the generated protocol through the chat interface.
3. Once the user is satisfied, the Main Agent invokes the Lab Knowledge Agent to save the final protocol as a new page in Confluence.

To supplement the general-purpose LLM with proteomics expertise, we optimized the Protocol Agent instructions. This multi-part prompt begins by instructing the model to adopt an expert persona in the field (e.g., “You are Matthias Mann”). It then provides contextual knowledge through domain-specific background information, such as images of laboratory equipment and detailed text descriptions of these (e.g., “An UltraSource from Bruker looks like a black spheroid housing with a thick white corrugated tube attached”). Furthermore, the prompt includes a clear task definition, which specifies the output for instance as a “Nature-style” protocol with standardized sections like abstract, materials, a step-by-step procedure and expected results.

To guide its reasoning, the prompt is formatted as chain-of-thought instructions, directing the model to first transcribe observations from the video with timestamps and then convert these observations into the structured protocol format. Finally, the prompt is grounded with few-shot examples, including a diverse set of complete video-protocol pairs that reflect possible user inputs, such as recordings from both a wearable camera and a screen. We found that our optimized prompt led to protocols that integrated both visual and verbal cues. In contrast, simple prompts (e.g., “Your task is to analyze the provided video and to convert it into a Nature-style protocol”) resulted in protocols that reflected primarily the spoken content.

##### Scenario C: Video analysis to detect errors

This workflow is triggered when a researcher submits an experimental video with a prompt like, “Generate lab notes from this video and check for mistakes.” (Appendix Figure S4). This requires a researcher to record themselves while performing an experiment.

1. The Main Agent delegates the initial analysis to the Lab Note Agent, which uploads the video to Google Cloud Storage, followed by video analysis to generate a summary of the procedure performed.
2. The Main Agent then invokes the Lab Knowledge Agent to search the lab’s Confluence for written protocols that match the video summary.
3. The agent suggests the best match to the user and asks for confirmation that this was the correct protocol.
4. Once confirmed, the Lab Knowledge Agent retrieves the full content of the correct protocol.
5. The Main Agent passes both the video and the baseline protocol to the Lab Note Agent for a detailed comparison. The agent systematically compares each step, flagging deviations such as omissions, additions, incorrect order, or procedural errors.
6. The agent outputs lab notes where potential errors are flagged for review.
7. After the researcher finalizes the notes, the Lab Knowledge Agent saves them to Confluence, preserving an accurate record of the experiment.

To detect errors during a lab procedure, we instructed the model to compare a written protocol with a video of the experiment. Our prompt directs the model to act as an expert, “Professor Matthias Mann”, and provides it with background documents on proteomics. The main task is to compare the video and the protocol, marking each step in the written protocol with a specific label: Correct, Omitted, Error, Added, or Deviation. This task follows an optimized chain-of-thought logic. The model begins by understanding the original written protocol and then goes through the video to create a timestamped log of all actions and spoken words. Next, it matches each event in the video log to the corresponding step in the protocol. For every potential mismatch it finds, the model re-examins the relevant video segment. After confirming all differences, the model generates the final annotated lab notes. The prompt also includes one complete example - a written protocol, a video, and the correct final output – to guide the task execution.

This error detection is a binary task as a step is either correct or wrong. Therefore, we adjusted the model’s temperature to a low value of 0.2, which makes the output more consistent. We avoided a temperature of 0, which often led to repetitive outputs.

#### Agent benchmarking

##### Benchmarking of protocol generation

We compiled a benchmark dataset of ten protocols recordings with a wearable camera (GoPro HERO8 black) covering a range of activities, from standard wet-lab procedures (e.g., pipetting) to interactions with specialized equipment (e.g., modifying MS ion sources) and software (e.g., instrument calibration) (Appendix Figure S3). These protocols were adapted from our recently published Nature Protocol about operating the mass spectrometer timsTOF from Bruker featuring the use of IonOpticks columns, Evotips that contain samples and Evosep as liquid chromatography system (Skowronek *et al*, 2025).

For automated evaluation, we used an ‘LLM-as-a-judge’ approach (Zheng *et al*, 2023), where a separate model compared the AI-generated protocols with manually created ground-truth, section by section and step by step. The judge rated each protocol on a 1-5 (5: very good) scale across five criteria: Completeness, technical accuracy, logical flow, safety instructions, and formatting. To produce the final overall quality score, a mean score was first calculated for each criterion across all sections and steps; these mean scores were then averaged.

##### Benchmarking of error detection

Using the ten protocols, we generated a benchmark dataset of 28 videos in which an expert scientist performed the procedures. Most videos contained intentionally introduced errors, while some were error-free controls. In total, the dataset comprised 421 steps, including 70 deliberate errors.

To evaluate performance, we compared the AI-generated lab notes against manually annotated ground-truth notes for each video. These ground-truth notes specified whether each step was correct or incorrect. Every step of the AI-generated lab notes was classified as a true positive, true negative, false positive, or false negative, which formed the basis for calculating accuracy, precision, and recall. For a quantitatively analysis of the agent’s strengths and limitations, errors were also categorized by their nature (omission, procedural error, added step, deviation of step order) and the perceptual skill required for detection (lab knowledge, proteomics knowledge, spatial resolution, spatial recognition, fast action recognition). This allowed us to calculate recall for each sub-category.

